# Long read mitochondrial genome sequencing using Cas9-guided adaptor ligation

**DOI:** 10.1101/2022.02.23.480720

**Authors:** Amy R. Vandiver, Brittany Pielstick, Timothy Gilpatrick, Austin N. Hoang, Hillary J. Vernon, Jonathan Wanagat, Winston Timp

**Author notes:** These authors contributed equally (Amy R. Vandiver, Brittany Pielstick). To whom correspondence should be addressed, Johns Hopkins University, 3400 N. Charles St, Clark 102A, Baltimore, MD 21218, University of California, Los Angeles, Los Angeles, CA 90095.

## Abstract

The mitochondrial genome (mtDNA) is an important source of disease-causing genetic variability, but existing sequencing methods limit understanding, precluding phased measurement of mutations and clear detection of large sporadic deletions. We adapted a method for amplification-free sequence enrichment using Cas9 cleavage to obtain full length nanopore reads of mtDNA. We then utilized the long reads to phase mutations in a patient with an mtDNA-linked syndrome and demonstrated that this method can map age-induced mtDNA deletions. We believe this method will offer deeper insight into our understanding of mtDNA variation.

## INTRODUCTION

Mitochondria are classically considered the power plants of eukaryotic cells, generating up to 90% of cellular ATP (Harris and Das). There is increasing evidence that mitochondria also play a larger role in regulation of cell behavior, including cellular identity, gene expression, cell death, and cellular senescence through modulation of the cellular metabolome (Antico Arciuch et al., 2012; Chakrabarty and Chandel, 2021; Picard et al., 2014; Sun et al., 2016). Most human cells contain multiple mitochondria, and each mitochondrion contains multiple copies of a 16,569 bp closed-circular genome which encodes 13 mRNAs, 22 tRNAs, and 2 rRNAs. The genes encoded are essential for mitochondrial function and interact with up to 1500 mitochondrial proteins encoded in the nuclear genome to form the electron transport chain (Anderson et al., 1981; Andrews et al., 1999).

While genetic variation within the nuclear genome is actively investigated and characterized, the patterns and extent of variation within the mitochondrial genome are relatively overlooked. Unlike the nuclear genome, mtDNA replicates independently from the cell cycle with a mitochondrial-specific, nuclear-encoded DNA polymerase. While mtDNA repair enzymes are present within the mitochondria, mtDNA accumulates polymorphisms at 10 to 100 times higher rate than nuclear DNA, likely due to differences in the function of the polymerase and repair machinery (Allio et al., 2017; Ludwig et al., 2019; Zheng et al., 2006). Analysis of cancer genomes indicates that the nuclear genome may influence the sites prone to variation within mtDNA (Hopkins et al., 2017).

Variants within mtDNA can be found in all mtDNA within a cell (homoplasmy) or in a subset of mtDNAs (heteroplasmy). Once present, specific polymorphisms have been observed to expand in a clonal manner, particularly in mitotic tissues (Nekhaeva et al., 2002). The relationship between heteroplasmic variants within a cell is not well understood due to the limits of current sequencing technology. Recombination of mtDNA may increase the diversity of genomes within a cell (Sato et al., 2005). Apart from point mutations, large deletions in mtDNA, ranging from 4-10 kbp, have been observed to accumulate with age, particularly in post-mitotic tissues (Herbst et al., 2021; Meissner et al., 2008). The mechanism of deletion formation remains unclear but may be linked to strand-slippage during mtDNA replication or repair of double-strand DNA breaks (Nekhaeva et al., 2002).

Deleterious mtDNA variants have been demonstrated to have different effects on cellular function depending on the specific variant, level of heteroplasmy, and metabolic demands of the cell type (Durham et al., 2007; Picard et al., 2014). Mitochondrial dysfunction is linked to a broad spectrum of pathology, ranging from maternally-inherited syndromes to oncogenesis and aging across many different cell and tissue types (Basu et al., 2020; Stewart et al., 2015; Yuan et al., 2020). A high heteroplasmic burden of specific single nucleotide polymorphisms or deletions is associated with multiple maternally-inherited diseases, including Leber’s hereditary optic neuropathy (LHON), mitochondrial encephalopathy, lactic acidosis and stroke like episodes (MELAS) and chronic progressive external ophthalmoplegia (CPEO), while lower heteroplasmy of many variants has been associated with many common age-related pathologies (Auré et al., 2007; Chomyn et al., 1992; Kellar-Wood et al., 1994; Sevini et al., 2014). Mitochondrial DNA deletions have been specifically linked to aging phenotypes in skeletal muscle (Herbst et al., 2021, 2016; Vermulst et al., 2008).

Though next generation sequencing has proved valuable in identifying mitochondrial sequence variation, there are limitations to these data. Single nucleotide variant calling with standard short-read sequencing protocols is limited by the high coverage needed to accurately classify heteroplasmy across a range of possible values. This is further complicated by the presence of nuclear-embedded mitochondrial sequences (NUMTs), fragments of mtDNA transferred to the nuclear genome during evolution. These range in size from 334 to 8798 bp, averaging 240 bp (Albayrak et al., 2016). These can be misaligned as mtDNA from short-read sequencing data and erroneously called as heteroplasmic mtDNA (Schon et al., 2012). Prior work has attempted to mitigate these concerns by enriching for mtDNA prior to sequencing through use of long-range mitochondrial PCR, exonuclease digestion of linear DNA or rolling circle amplification (Palculict et al., 2016; Yao et al., 2019). However, the use of PCR in these methods has the potential to bias representation, particularly against mtDNA containing structural alterations. Further, methods reliant on short read sequencing and PCR have limited use for understanding the relationship between mtDNA variants within a cell, as they have limited capacity to determine which variants co-segregate within a single mitochondrial genome (Alkanaq et al., 2019).

Here, we use a recently developed Cas9 based enrichment method (nanopore Cas9-targeted sequencing; nCATS) on the Oxford nanopore long-read platform(Gilpatrick et al., 2020) to address the existing limitations in mtDNA sequencing. We demonstrate that mtDNA nCATS is effective for mtDNA enrichment and generates full-length mtDNA sequences. Using a well-characterized lymphoblast cell line, we demonstrate that nCATS enrichment generates single nucleotide polymorphism (SNP) data equivalent to short read sequencing. Using human muscle samples, we demonstrate that nCATS enrichment allows for localization of age-induced mtDNA structural variants. Finally, using peripheral blood of a patient with an mtDNA-associated syndrome, we determine the patterns of co-segregation of potential pathogenic mtDNA variants.

## MATERIALS AND METHODS

### nCATS Sequencing

nCATS sequencing was performed in accordance with previously described methods(Gilpatrick et al., 2020). Briefly: custom guide RNA, sequence: ACCCCTACGCATTTATATAG was designed using Benchling guideRNA design tool and checked for off target mtDNA sites using IDT design tool. Ribonucleoprotein complex assembly was performed by mixing crRNA and tracrRNA (IDT, cat no 1072532) at a concentration of 10 uM each. Duplexes were formed through denaturation and cooling, then combined with HiFi Cas9 Nuclease V3 (IDT, cat 1081060) in 1X CutSmart Buffer (NEB). Next, 3 ug of input DNA was dephosphorylated with Quick CIP (New England Biolabs, cat no M0508). Following enzyme inactivation with alkaline phosphatase, 10 uL of 333 nM Cas9-gRNA complex was added to the sample. After Cas9 cleavage, sequencing adaptors were ligated to DNA ends using the Oxford Nanopore ligation kit (LSK-109). Samples were run on a MinION (v9.4.1) flow cell using the GridION sequencer. Sequencing runs were operated using the MinKNOW software (v19.2.2).

### Illumina mtDNA sequencing

Mitochondrial DNA was amplified using the REPLI-g Mitochondrial DNA kit (Qiagen, cat no 151023) according to manufacturer’s instructions. Illumina library prep was performed using Nextera DNA Flex Library Preparation kit (cat no. 20018704) and sequenced on the iSeq100 (Illumina).

### GM12878 Cell culture and DNA prep

CEPH/UTAH Pedigree 1463 (GM12878) cells were obtained from Coriell Institute and cultured according to recommended protocols. Briefly, cells were grown in high-glucose RPMI medium supplemented with 10% fetal calf serum, penicillin-streptomycin antibiotics and L-glutamine. Cells were maintained at 37°C at 5% CO_2_. DNA was extracted using the MasterPure kit (Lucigen, cat no MC85200).

### Muscle sample collection and DNA prep

De-identified muscle biopsy specimens were collected as part of a VA Merit Award, “Testosterone, inflammation and metabolic risk in older Veterans” and NIH R01K090406. Use of the human specimens for this study was approved by the UCLA Health Institutional Review Board (Protocol #18-001547). Core muscle biopsies were taken from quad muscle, flash frozen and stored at −80°C prior to DNA extraction. Tissue samples were powdered under liquid nitrogen using a mortar and pestle. Approximately 25 mg of powdered muscle was used for DNA isolation, performed by proteinase K and RNase A digestion followed by phenol-chloroform extraction and ethanol precipitation, as previously described (Herbst et al., 2021).

### Blood sample collection and DNA prep

Whole blood was collected with patient consent through the Mendel Project which permits sharing de-identified clinical data, database submissions and publications. DNA isolation was performed using Circulomics Nanobind CBB big DNA kit (cells, Bacteria, Blood) (NB-900-001-01).

### Data Analysis

#### Alignment

Nanopore sequencing runs were basecalled using guppy (v5.0.11) to generate FASTQ sequencing reads. Alignment of nanopore reads was performed using minimap2 (2.17) (Li, 2018). Data was aligned to a rotated version of the revised Cambridge mitochondrial genome sequence, for which base pairs 1 through 1547 were shifted to the end of the genome to account for location of Cas9 cleavage. Illumina sequencing data was aligned to the same modified mitochondrial genome using BWA (v0.7.17) (Li and Durbin, 2009).

#### Variant calling

Variant calling in nanopore data was performed using bcftools (v1.9) (Li, 2018) call function with ploidy set to 1. Calls were filtered to include only variants with coverage >10% of total chromosome M reads. Variant calling in Illumina data was performed using FreeBayes (v1.3.2). High confidence variants were identified through use of vcf filter to identify variants for which alternative allele reads were found on both strands. An insertion called at the locations of an artificially introduced N in the mitochondrial reference genome at bp 3106 (Andrews et al., 1999) was filtered from final variant lists.

#### Deletion finding

Deletions were identified on a single read level from aligned data by parsing the CIGAR string in GenomicAlignments (v1.26.0) to identify contiguous deleted bases greater than 100 bp.

#### Variant phasing

Phasing of variants in blood sequencing data was performed by querying the PySam (v0.15.3) pileup of all reads with no quality filter to identify the base pair at each site of interest for all reads covering both sites.

Analysis code is available at https://github.com/amyruthvandiver/Cas9_Mitochondria

## RESULTS AND DISCUSSION

### Cas9-based enrichment for targeted mitochondrial genome sequencing on Oxford Nanopore

In order to address the current limitations of next generation sequencing approaches for understanding the mitochondrial genome, we adapted nCATS, a Cas9 based long-read enrichment method to enrich mtDNA. This method capitalizes on the selective ligation of nanopore sequencing adaptors to enrich genetic regions of interest. Samples are first de-phosphorylated at all free DNA ends, then Cas9 cleavage creates blunt ends with retained 5’ phosphorylated ends (Gilpatrick et al., 2020). We predicted that the use of a guide RNA specific to mtDNA would linearize mtDNA and lead to dramatic enrichment of mtDNA in nanopore sequencing data (Figure 1A). To test this approach, a guide RNA was designed to cut mtDNA once within the sequence encoding ribosomal RNA1 at bp 1547. This site was chosen due to the low frequency of reported single nucleotide variants and deletions affecting this region in the MITOMAP database (Brandon et al., 2005) (Figure 1C) and the low rate of off-target cuts against the reference human genome.

**Figure 1:**
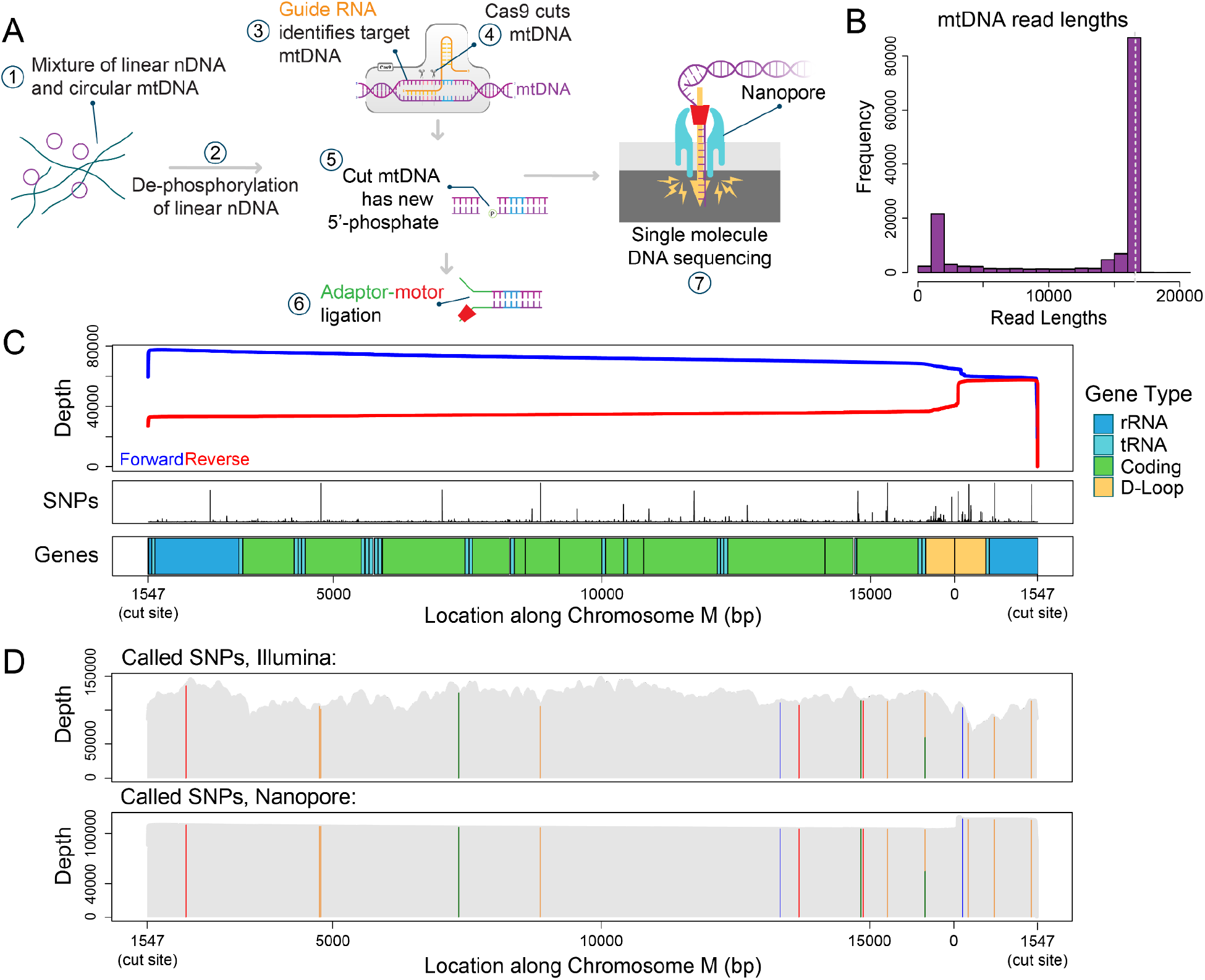
Targeted mtDNA sequencing in GM12878 cells. **A)** Adaptation of nCATS protocol to selectively sequence mtDNA. Beginning with total DNA, free ends are dephosphorylated, followed by linearization and introduction of free phosphorylated ends in mtDNA by Cas9 cleavage. Adaptor sequences complexed with motor protein are selectively ligated to available phosphorylated ends and the targeted regions are sequenced. **B)** Histogram depicting length distribution of reads aligning to chromosome M in nanopore sequencing of GM12878 cells. Dashed line indicating full length mtDNA. **C)** Top panel: Sequencing depth for forward (blue) strand and reverse (red) strand across chromosome M. Middle panel: SNP frequency in the MitoMap database. Bottom panel: Annotation of chromosome M. **D)** SNP calling in GM12878 cell line. Top panel: Location of SNPs identified in GIAB Illumina data, sequencing depth in grey. Bottom panel: Location of SNPs identified in nanopore data, sequencing depth in grey.

For initial validation of this method, we used the well characterized lymphoblast line, GM12878. From this cell line, we generated 193,043 reads, of which 142,373 (73.8%) aligned to the revised Cambridge mitochondrial reference genome (Andrews et al., 1999), rotated to begin at the Cas9 cut site (bp 1547). The median length of primary alignments was 16,458 bp, with 66% of chrM reads greater than 15 kb (Figure 1B, Supplementary Table 1); note the mtDNA reference is 16,569 bp long. Notably we observed a significant bias towards forward strand reads versus reverse strand reads (Figure 1C), this has previously been observed in Cas9-based enrichment (Gilpatrick et al., 2020; Wallace et al., 2021), likely because of how residual Cas9 blocks ligation of the adaptor to one side of the cut.

Examination of the lengths of chrM reads showed a secondary peak between 1450 and 1550 bp, accounting for 11% of chrM reads (Figure 1B). Of these, 85% align with a start between bp 56 and 76 and end at the Cas9 cut site. This region falls within the mitochondrial D-loop near the origin of replication, suggesting the structure in this region leads to Cas9-independent fragmentation. While our reported percent of mtDNA sequences is lower than the previously reported highest mtDNA percent achieved through exonuclease digestion and rolling circle amplification (Yao et al., 2019) it is significantly higher than other amplification-free methods of enrichment (Gould et al., 2015) and provides dramatically increased read length from all NGS-based methods.

To determine the utility of this method for genotyping mtDNA, we compared our nanopore data to publicly available high coverage Illumina sequencing of GM12878 cells from the Genome in a Bottle (GIAB) project (Zook et al., 2016). For nanopore data, high confidence single nucleotide polymorphisms (SNPs) were called using bcftools and filtered to include only variants covered by >10% of reads and detected on both strands. For Illumina data, SNPs were called using FreeBayes, and then filtered to correct for strand bias. In both data sets, 15 identical single nucleotide polymorphisms were identified, of which 14 were homoplasmic. The single heteroplasmic SNP at bp 6203, was identified in 53% of reads in nanopore sequencing data and 48% of reads in Illumina data (Figure 1D, Supplementary Table 2).

### Phasing of mtDNA point mutations in long-read sequencing

The full-length mtDNA reads from nanopore sequencing allow us to investigate co-segregation of mitochondrial variants within mtDNA molecules; this is not possible with short read sequencing. As a proof of principle, we used blood sample DNA from a patient with MELAS syndrome, presenting with kidney failure and neuronal deficits. Prior whole exome sequencing of this patient had identified two putative deleterious G to A mutations in the mitochondrial genome at bps 1642, within the valine tRNA, and 13,513, within the coding region for *ND5*, both present in 8% of sequences. Both of these variants have been previously reported in association with mitochondrial disease. The G1642A variant was identified in 2 patients with MELAS (de Coo et al., 1998; Taylor et al., 1996). The G13513A variant has been reported in association with a variety of clinical presentations including MELAS, Leigh disease and isolated optic neuropathy (Shanske et al., 2008; Sun et al., 2021). Given the patient’s clinical presentation, it was concluded that the G13513A variant was likely responsible for her disease, with the G1642A variant possibly contributory, however the relationship between the variants is unknown. Determining whether the variants reside on the same or different mitochondrial genomes is critical to understanding whether the patient’s siblings and offspring are at risk for a single disease entity or two independent disease entities.

To explore the co-segregation of these two variants, we extracted DNA from peripheral blood from the patient of interest. The presence of the previously identified mutations was validated by Illumina sequencing of mtDNA amplified by rolling circle amplification, yielding 51,718 reads aligned to mtDNA, with 413x (chrM:1642) and 300x (chrM:13,513) coverage at the sites of interest. The genotype at each site of interest was measured using bcftools; we found 8% G1642A variants and 6% G13513A variants (Figure 2A-B). We then performed the validated nCATS-mtDNA to generate full-length mtDNA reads. From this sample, we generated only 7,042 total reads, which we attribute to the relative difficulty of extracting high quality mtDNA from this sample. However, 2,218 reads aligned to the reference mitochondrial genome with an average length of 11,673 bp. Initial examination of the sites of interest using bcftools revealed similar levels of G to A heteroplasmy as identified in the Illumina data, with 7.6% 1642A and 7.2% 13531A (Figure 2A-B).

**Figure 2:**
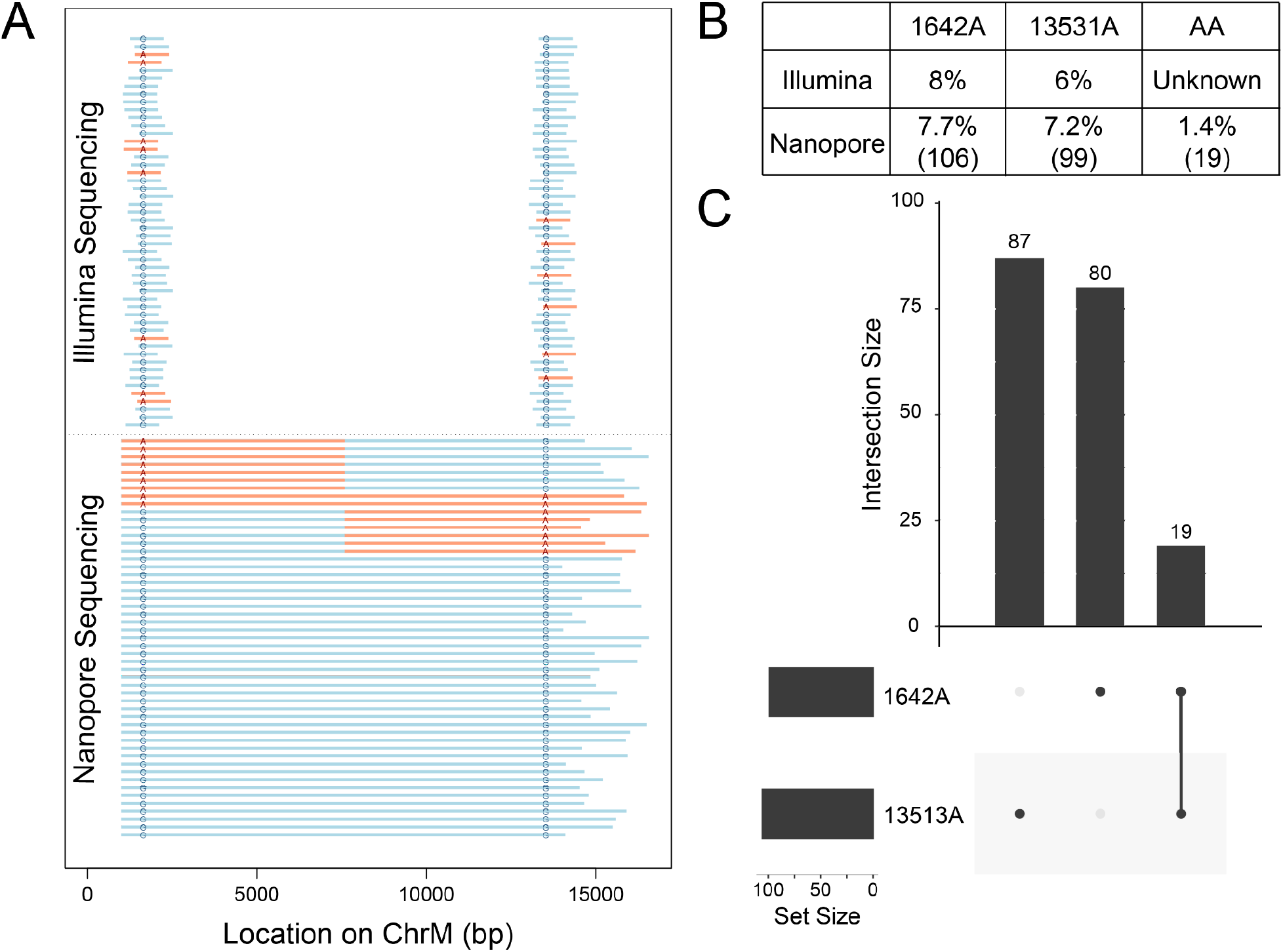
Phasing of mtDNA point mutations using targeted long read sequencing. **A)** Graphical depiction of representative Illumina and nanopore sequencing data with highlighted genotypes at the sites of interest. Representative reads with SNP shown in red, reference genotype in blue. For ease of view, 10% of mutant reads are shown, while 2.5% of double reference reads are shown. **B)** Table denoting percentage of reads of each genotype detected in each type of sequencing data. For nanopore data, number of reads of each genotype shown in parentheses. **C)** Upset plot showing co-segregation of G to A SNPs in nanopore sequencing data.

A total of 1,369 reads simultaneously spanned both SNP loci; using pysam to query the genotypes at both sites for each of these reads, we identified four combinations of mtDNA: 1,183 (86%) with neither variant, ‘GG’, 87 reads (6.4%) with only the first variant ‘AG’, 80 reads (5.8%) with only the second variant ‘GA’, and 19 reads (1.4%) with both variants, ‘AA’ (Figure 2, Supplementary Figure 1). To determine if nanopore sequencing error could account for the presence of the low frequency double variant population, we repeated this analysis on all reads from the GM12878 cell line and six human muscle samples. No control sample showed heteroplasmy of either 1642A or 13531A > 0.5% of reads and no sample showed ‘AA’ in more than 0.01% of reads, suggesting that the levels of variation observed in the MELAS patient’s blood sample are unlikely attributable to sequencing error alone (Supplementary Table 3).

The identification of four different populations - the reference, either single variant, or both variants, was a surprising finding. Finding the two variants both alone and together could indicate that recombination events are present as has been previously hypothesized (Sato et al., 2005) or that these specific sites are prone to somatically-acquired variation in this specific nuclear background, as hypothesized to occur in cancer evolution (Hopkins et al., 2017). The relatively low levels of heteroplasmy of the SNPs of interest are consistent with prior reports of selection against pathogenic mtDNA in blood samples of MELAS patients, even while heteroplasmy increases in disease related tissue (Rajasimha et al., 2008). It is thus possible that higher levels of heteroplasmy and a difference in distribution of mtDNA populations may be observed when other tissues from this patient are analyzed.

### Identification of structural variants in mtDNA of aged muscle samples

Large deletions within the mitochondrial genome have been repeatedly observed in diverse settings including cancer, sun exposure and aging (Berneburg and Krutmann, 1998; Meissner et al., 2008; Yusoff et al., 2019); however, the breadth and localization of these changes have been challenging to resolve using short read sequencing data (Bosworth et al., 2017). To determine the utility of our method for resolving structural changes, we utilized skeletal muscle, a tissue in which a 98-fold increase in mtDNA deletions has been demonstrated over the human lifespan using digital PCR (Herbst et al., 2021). For our initial test, DNA was extracted from six core biopsy samples of quadriceps muscle obtained from donors ranging from 20 years to 86 years of age. From each sample, mtDNA was sequenced using our nCATS-mtDNA protocol. Application of this method to muscle DNA demonstrated effective mtDNA enrichment, with 76-97% aligning to chromosome M, though sequencing varied substantially in output, with chrM reads ranging from 4071 to 145,127. Mean mitochondrial read length ranged from 3888-12757 bp (Supplementary Table 1).

To identify deletions within the obtained reads, the CIGAR strand was queried from the alignments to identify the length, start and end of single read deletions greater than 100 bp. Examination of these identified deletions revealed a large number of deletions <2 kbp in both younger and older samples (Figure 3A, Supplementary Table 4). Notably however, deletions larger than 2kbp are predominantly (91%) found in samples from donors >60 years old (Figure 3B, Supplementary Table 4). All but two of the deletions >2kbp were found within the mitochondrial major arc region spanning bp 407-5747 (Figure 3C), consistent with the distribution of previously reported deletions found with NGS, long extension PCR, and single cell analyses (Brandon et al.)(Bua et al., 2006). The fraction of reads with long deletions ranged from 7 x 10^-5^ to 5 x 10^-4^ in the younger three samples to a higher level of 7 x 10^-4^ to 3 x 10^-3^ in the older three samples. The human mitochondrial “common deletion,” a deletion spanning tandem repeat sequences at bp 8470-13477 which is associated with Kearns-Sayre syndrome and frequently observed deletion in studies of aged tissue (Meissner et al., 2008), was observed 4 times in the younger samples and 10 times in the older samples, accounting for 4% of the total deletions identified. A BED file detailing each identified deletion is provided as Supplementary Data 1.

**Figure 3:**
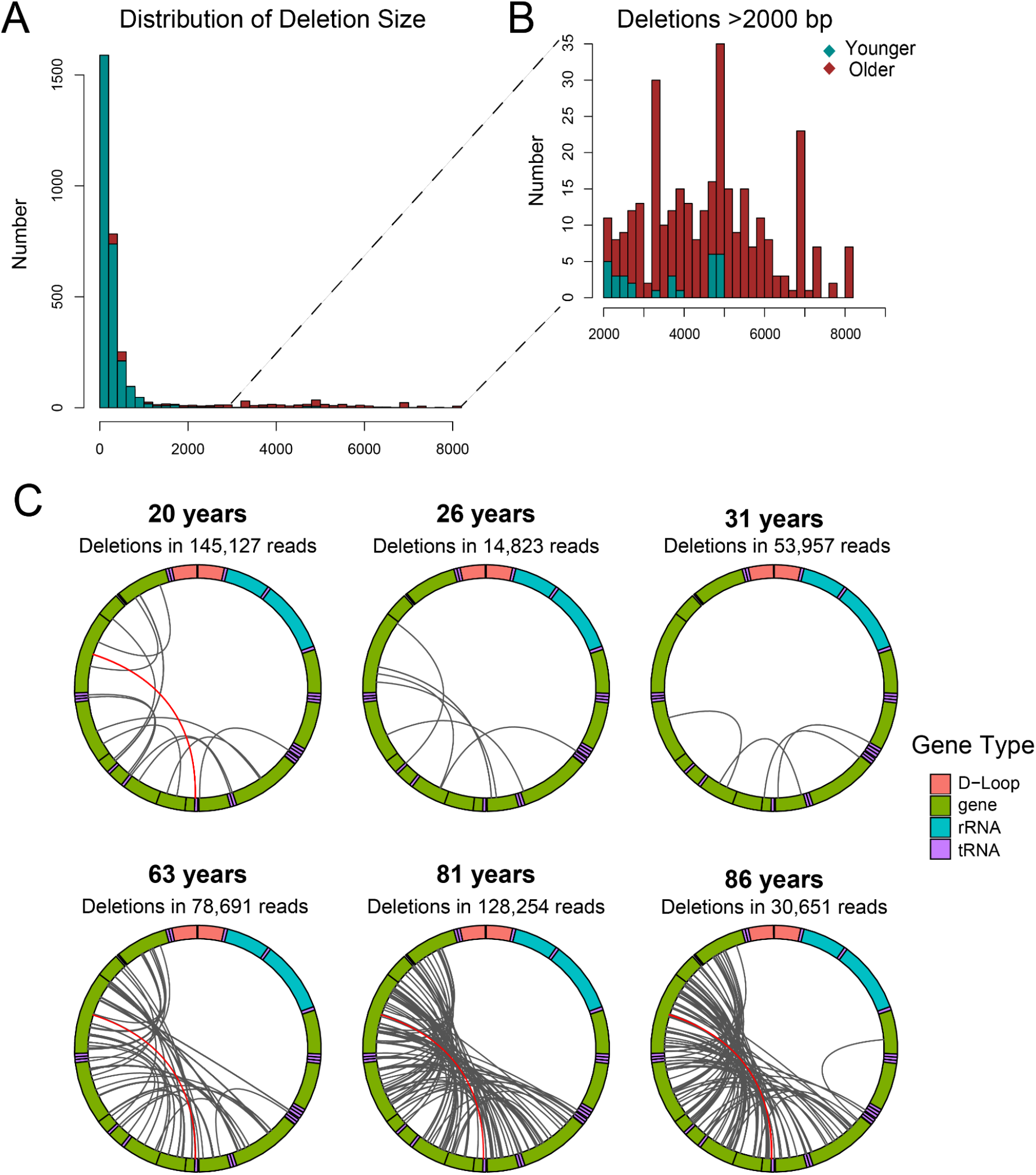
Identification of mtDNA deletions using targeted nanopore sequencing. **A)** Distribution of mtDNA deletion lengths for deletions >100 bp in samples from donors <35 years (green) and samples from donors >60 years (red). **B)** Distribution of mtDNA deletion lengths for deletions >2000 bp in samples from donors <35 years (green) and samples from donors >60 years (red). **C)** Localization of all deletions >2000 kbp identified per sample. Mitochondrial genome depicted in clockwise orientation with lines linking the start and end of each identified deletion event. The human mitochondrial “common deletion” is highlighted in red where identified.

## CONCLUSIONS

Here we demonstrate the use of targeted nanopore sequencing using nCATS to obtain high coverage, full-length sequencing of the mitochondrial genome. This method provides consistent enrichment of mtDNA in a cell line, human blood, and human muscle tissue samples. Our data shows the utility of full-length characterization for identifying both cosegregation of point mutations and of structural variants, primarily deletions, within heteroplasmic mtDNA.

## DATA AVAILABILITY

GM12878 targeted data are available at SRA under Bioproject ID PRJNA809571 (https://www.ncbi.nlm.nih.gov/bioproject/809571).

## ACKNOWLEDGEMENTS

AV has received support from NIH/NIAMS T32 AR071307 and The Dermatology Foundation. This work was funded in part from NIH R01HG009190 (WT, BP, TG), NIH T32 GM007445 (BP), and NIH T32 GM136577 and (R56AG060880, R01AG055518, and K02AG059847) to JW.

## AUTHOR CONTRIBUTIONS

BP, AV, and WT constructed the study. BP and TG performed the experiments. BP, AV, and WT analyzed the data. AH and JW provided primary muscle tissue and generated muscle DNA. HV provided primary blood samples and extracted blood DNA. AV, BP, and WT wrote the manuscript.

## SUPPLEMENTARY INFORMATION

**Supplementary Figure 1:** Co-localization of G to A variants in single mtDNA reads from MELAS patient. Shown are integrated genome viewer snapshots depicting the called based at positions 1642 and 13513 for a subset of reads containing the 1642A variant and a subset of reads containing the 13513A variant.

**Supplementary Data 1:** BED file containing the locations of all deletions identified.

**Supplementary Table 1:** Summary of all sequencing runs, including sample name, read count, mtDNA enrichment and summary statistics of mtDNA reads.

**Supplementary Table 2:** SNPs identified in Illumina and Nanopore data from GM12878 cells.

**Supplementary Table 3:** Raw counts and frequencies of 1642A and 13513A SNPs detected in all samples.

**Supplementary Table 4:** Counts and read fraction of deletions <2 kbp and >2 kbp detected in each muscle sample.

