## Supplementary Figures for "Long read mitochondrial genome sequencing using Cas9-guided adaptor ligation"

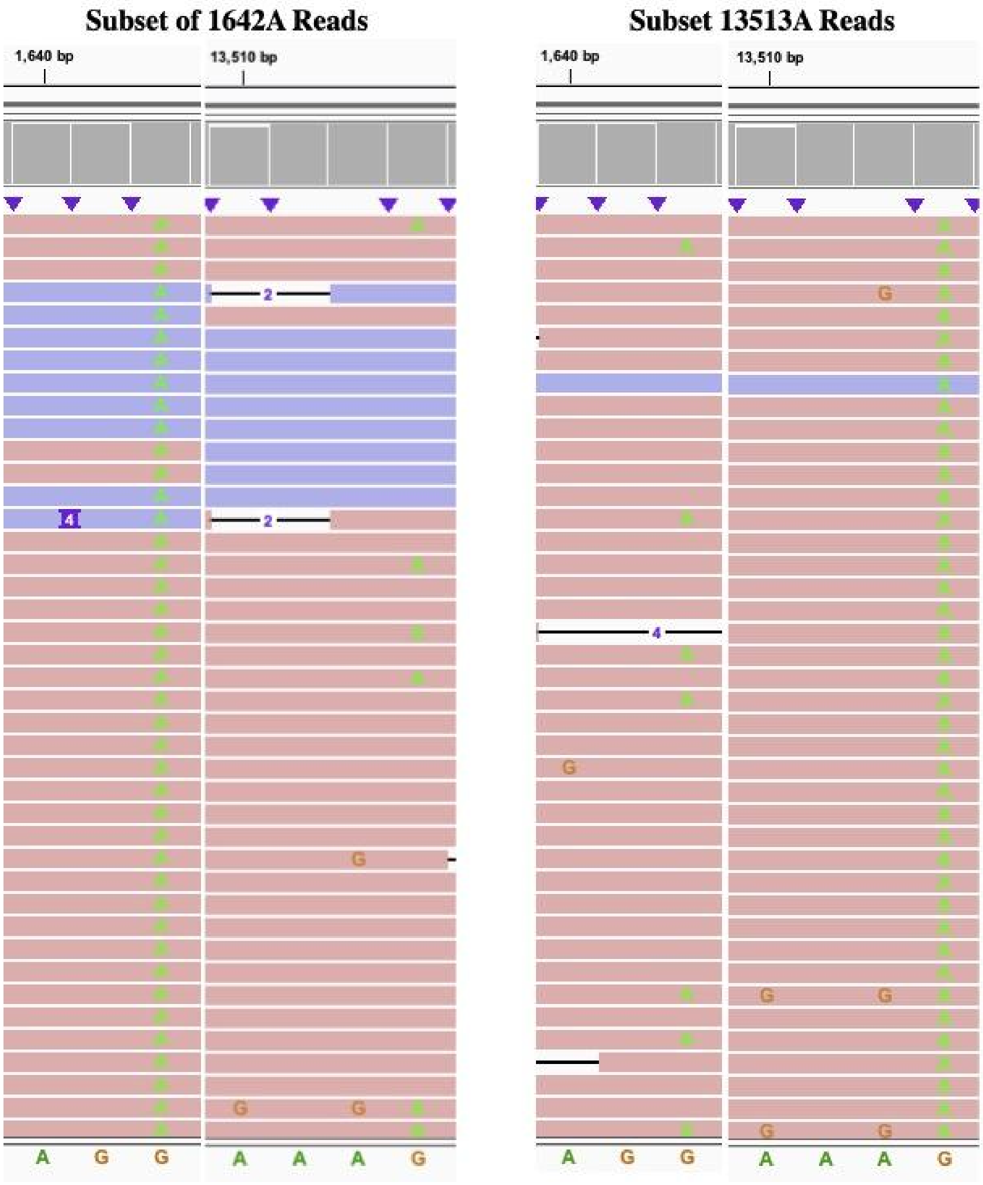


***Supplementary Figure 1: Co-localization of G to A variants in single mtDNA reads from MELAS patient.*** *Shown are Integrated Genome Viewer (IGV) snapshots depicting the called bases at position 1642 and 13,513 for a subset of reads containing the 1642A variant and a subset of reads containing the 13513A variant.*
